# The base media formulation impacts the efficiency of ex-vivo erythropoiesis of primary human hematopoietic stem cells

**DOI:** 10.1101/2023.12.24.573281

**Authors:** Nana Ansuah Peterson, Elizabeth S. Egan

## Abstract

During the COVID-19 pandemic, there was a disruption in the supply of the widely used erythroid differentiation media product, Iscove’s Modified Dulbecco’s Medium, manufactured by Biochrom AG. IMDM is a critical component of ex vivo erythropoiesis protocols used to generate genetically modified red blood cells for the study of malaria host factors. Therefore, we set out to identify the best alternative IMDM product for efficient erythroid differentiation of hematopoietic stem cells into enucleated red blood cells for use in our research. We tested other IMDM products, including I2911 (from Millipore Sigma), P04-20450 (from Pan Biotech) and EP-CM-L0216 (from ElabScience Technologies) which are all marketed specifically as replacements for the Biochrom AG product. We found that while FG0465, I2911, and P04-20450 were all sufficient for ex-vivo erythropoiesis, FG0465 IMDM was superior to other products tested in supporting maximal erythroid cell proliferation and enucleation.

## Introduction

Ex-vivo erythropoiesis has implications for patient care as well as translational and basic science research. Large-scale production of human erythrocytes in vitro is currently being explored for potential clinical applications such as blood transfusions; the first clinical trial for the transfusion of red blood cells grown in vitro is currently underway^1^. As the obligate host cell for the asexual replication of *Plasmodium falciparum* parasites, erythrocytes are critical to the culture and study of malaria parasites. Since mature human erythrocytes are terminally differentiated and lack nuclei, making them genetically intractable, recent advances in the ex-vivo differentiation of stem cells to enucleated reticulocytes and erythrocytes is enabling innovative research into the host genetic determinants of malaria^2-4^. Genome engineering to generate genetically altered erythrocytes must begin with CD34^+^ hematopoietic stem/progenitor cells (HSPCs), which can be genetically manipulated and differentiated down the erythropoietic lineage. In addition to malaria, such approaches are also foundational for research into the genetics of terminal erythroid differentiation, bone marrow failure syndromes such as Diamond-Blackfan Anemia, and hemoglobinopathies.

Ex-vivo erythropoiesis approaches have a long history, but recent progress has significantly advanced the reproducibility and efficiency of this process. In 2005, Giarratana et al published a protocol to generate erythroid precursors from primary human CD34^+^ HSPCs which proliferate, terminally differentiate and enucleate efficiently to produce substantive yields of mature erythrocytes^5^. Key to this protocol is erythroid differentiation medium (EDM) which is made using Iscove’s Modified Dulbecco’s Medium (IMDM) as the base media. IMDM is a derivative of Dulbecco’s culture medium modified to support the culture and differentiation of hematopoietic precursors by the addition of selenite, pyruvate, some additional amino acids, changing the concentration of several salts and the replacement of ferric nitrate with potassium nitrate^6,7^. To promote erythroid proliferation and differentiation, Giarratana and colleagues supplemented IMDM with glutamine, holotransferrin, insulin, heparin choay, serum, IL-3, hydrocortisone, erythropoietin (EPO) and stem cell factor (SCF)^5,8,9^. To facilitate enucleation, the differentiating cells were co-cultured on a murine stromal layer towards the end of the culture. In subsequent work, the same group showed that use of human plasma could obviate the need for co-culture on a stromal layer, still leading to highly efficient enucleation^10^.

The ex-vivo erythropoiesis protocol developed by Douay and Giarratana made use of IMDM specifically from the manufacturer Biochrom AG (catalog number F0465). Biochrom AG F0465 required supplementation with l-glutamine, but more recently this product became available with stabilized glutamine already added, and is currently sold by Sigma (catalog number FG0465). Many other recent studies involving ex-vivo erythropoiesis for the study of malaria host factors have similarly relied on F/G0465 IMDM^2,11,12^. While other manufacturers market IMDM products as being similar or identical to FG0465, to our knowledge their efficacy for supporting the proliferation and erythroid differentiation of primary human HSPCs has not been tested. With supply chain issues intermittently impacting the availability of FG0465, we sought to investigate the potential of other IMDM products to support ex-vivo erythropoiesis of primary HSPCs to enucleated red blood cells.

Here, we used a set of quantitative phenotypic assays to compare the efficiency of ex-vivo erythropoiesis of primary human HSPCs in 4 different IMDM products. Our results demonstrate that the efficiency of erythroid precursor cell proliferation and terminal differentiation to enucleated RBCs are dependent on the precise formulation of IMDM used for ex-vivo erythropoiesis.

### Methods Culture media

CD34^+^ primary human hematopoietic stem cells (HSPCs) isolated from human bone marrow were cultured in erythroid differentiation media (EDM) made from one of four different IMDM products supplemented with 330 μg/ml human holotransferrin (BBI Solutions), 10 μg/ml recombinant human insulin (Sigma), 2 IU/ml heparin choay (Affymetrix) and 5% human plasma (Octapharma). The four different IMDM formulations─ each described as containing sodium bicarbonate, D-glucose, L-alanyl-l-glutamine and phenol red─were FG0465 (Biochrom AG product currently supplied by Millipore Sigma), I2911 (manufactured and supplied by Millipore Sigma), P04-20450 (manufactured by Pan Biotech) and EP-CM-L0216 (manufactured by ElabScience Technologies). In some experiments, 4mM L-glutamine (Sigma) was added; EDM prepared from P04-20450 was always supplemented with L-glutamine whereas EP-CM-L0216 and I2911 were tested with and without L-glutamine (Table 1).

**Table 1:** IMDM +/-added L-Glutamine used in the different experiments.

|  | FG0465 | EP-CM-L0216 | EP-CM-L0216 +Glut | I2911 | I2911 +Glut | P04-20450 +Glut |
| --- | --- | --- | --- | --- | --- | --- |
| Experiment 0 |  |  |  | ✓ |  |  |
| Experiment 1 | ✓ | ✓ | ✓ | ✓ | ✓ |  |
| Experiment 2 | ✓ |  |  |  |  | ✓ |
| Experiment 3 | ✓ |  |  |  | ✓ | ✓ |

As previously described^3,8,12^, EDM was supplemented with the following cytokines and hormones: 5 ng/ml interleukin-3 (IL-3, R&D systems), 1 μM hydrocortisone (HC, StemCell Technologies), 100 ng/ml stem cell factor (SCF, R&D Systems) and 3 IU/ml erythropoietin (EPO, Amgen).

### Cell culture

3 × 10^5^ or 1 × 10^5^ CD34^+^ bone marrow HSPCs (StemCell Technologies) were thawed in FG0465 EDM supplemented with 2.5% plasma, and then split evenly among the different IMDM-based EDM preparations supplemented with human plasma, IL-3, HC, SCF and EPO, plated at ∼1 × 10^4^ cells /ml.

On day 4 of culture, the cells were washed in EDM supplemented with 2.5% plasma and plated in 5x the starting volume of EDM supplemented with plasma, IL-3, HC, SCF and EPO. On day 7-8, the cells were washed and plated at 2-5 × 10^5^ cells/ml in EDM supplemented with plasma, SCF and EPO. On day 11 of culture, the cells were washed in EDM and plated at 0.75-1 × 10^6^ cells/ ml in EDM supplemented with plasma and EPO. From day 12-13 of culture, cells were co-cultured on a murine stromal layer (MS-5) in EDM with plasma and EPO to facilitate enucleation. Cells were counted using a hematocytometer to estimate total cell numbers every 2-3 days throughout the culture.

### Microscopy

Cytospins were stained in May-Grünwald and Giemsa stains for morphological analyses. Images of cytospins were taken on a Keyence BZ-X700 microscope with a 60X magnification oil objective lens.

### Flow cytometry

Expression of cell surface proteins including transferrin receptor 1 (CD71), integrin alpha 4 (CD49d) and glycophorin A (GPA) was assessed on days 11-12 and 17-18 by flow cytometric analysis of immunostained cells. Cells were washed in 0.3% PBS/BSA and stained in 1:20 dilution of anti-GPA-APC-efluor780 clone HIR2 (Invitrogen) or 1:50 dilution of anti-GPA-FITC clone 2B7 (StemCell Technologies) or 1:10 dilution of anti-CD71-PE (Miltenyi) or 1:10 dilution of anti-CD49d-APC clone M218-24A9 (Miltenyi). Enucleation rates were assessed by flow cytometric analyses of cells stained in 1:1000 dilution of Vybrant DyeCycle Ruby Stain (Invitrogen) in RPMI-1640 (with 25 mM HEPES and 50 mg/L hypoxanthine). Samples were run on a MacsQuant flow cytometer (Miltenyi).

### Data Analysis

Flow cytometry data were analyzed in FlowJo version 10.8.1. Statistical analyses were performed, and graphs were generated in GraphPad prism version 9.3.1 for Windows.

## Results

### Erythroid cell proliferation is influenced by the IMDM base media

To determine the impact of the base IMDM formulation on the efficiency of ex-vivo erythropoiesis, we prepared EDM using four different commercially available IMDM products (FG0465, EP-CM-L0216, I2911, and P04-20450). While our prior work has shown that FG0465, which is marketed as containing stable glutamine, can support ex-vivo proliferation and differentiation of primary human hematopoietic stem cells without additional glutamine^12^, for the other base media formulations we tested conditions with and without glutamine supplementation.

As demonstrated previously^8,12^, EDM made with FG0465 IMDM was sufficient to support expansion of primary human hematopoietic stem cells, with >2000-fold cumulative fold change over 15 days of culture (Fig. 1). In contrast, CD34^+^ HSPCs incubated in EDM made with I2911 failed to proliferate unless the culture was supplemented with glutamine (Fig. 1). By day 7 of differentiation, very few to no viable cells could be seen in the I2911 EDM without glutamine, whereas additional supplementation of I2911-based EDM with glutamine enabled cell proliferation (Fig. 1 inset). However, the degree of erythroid cell expansion in I2911 EDM with glutamine (I2911+Glut EDM) was consistently less than in FG0465 (800-fold versus 2000-fold by day 15) (unpaired one-tailed student’s t-test, n= 2, p-value 0.0113) (Fig. 1). Similarly, EDM made with P04-20450 with additional glutamine supplementation was sufficient to sustain growth, but lower cell yields were observed compared to FG0465 EDM (unpaired one-tailed student’s t-test, n=2, p-value 0.0242) (Fig. 1). Strikingly, EDM made with EP-CM-L0216 was not sufficient to sustain the survival of primary human CD34^+^ HSPCs regardless of additional glutamine supplementation (Fig. 1).

**Figure 1:**
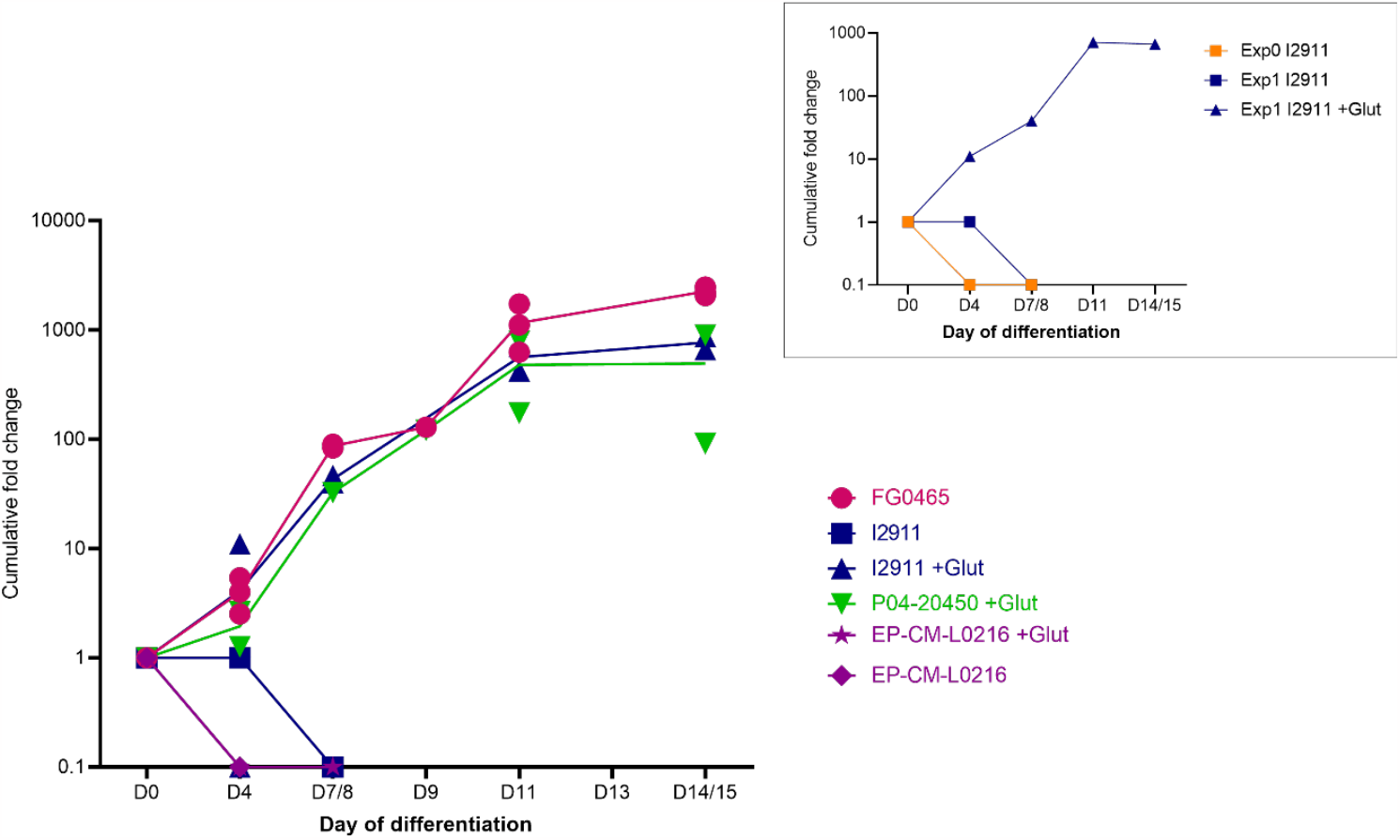
Erythroid cell expansion is higher in EDM made with FG0465 than in other IMDM tested. Cumulative fold change by day of differentiation for cells grown in EDM made from different IMDM with/ without added L-glutamine across different experiments. Solid lines connect the means of the replicates over the timecourse. FG0465 (n= 3), EP-CM-L0216 (n=1), EP-CM-L0216 + Glut (n=1), I2911 (n=1), I2911 + Glut (n=2), P04-20450 + Glut (n=2). The figure inset compares cumulative fold change for cells grown in I2911-EDM compared to I2911+Glut-EDM. Exp1 data points in inset are also displayed in the main figure.

### EDM made with FG0465, I2911 (+glut) and P04-20450 (+glut) are sufficient to support erythroid differentiation

We next sought to assess the capacity of primary human CD34^+^ HSPCs to differentiate down the erythroid lineage when cultured in EDM made with different IMDM base media formulations. To do this, we harvested the cells at specific timepoints during erythroid differentiation and used both microscopy and flow cytometry to assess their developmental progression. Erythroid precursors grown in EDM made from FG0465, I2911 supplemented with glutamine (I2911 +glut), and P04-20450 supplemented with glutamine (P04-20450+glut) progressed through erythroid developmental stages in a similar manner, as assessed by light microscopy (Fig. 2). On day 11, most cells appeared to be basophilic or polychromatic erythroblasts, with occasional orthochromatic erythroblasts observed. By day 18, most cells were enucleated reticulocytes, consistent with effective terminal differentiation. However, the cells cultured in I2911+glut-EDM or P04-20450+glut-EDM appeared to have a lower proportion of enucleated cRBCs on day 18-19 relative to those grown in EDM made with FG0465 (Fig. 2B).

**Figure 2:**
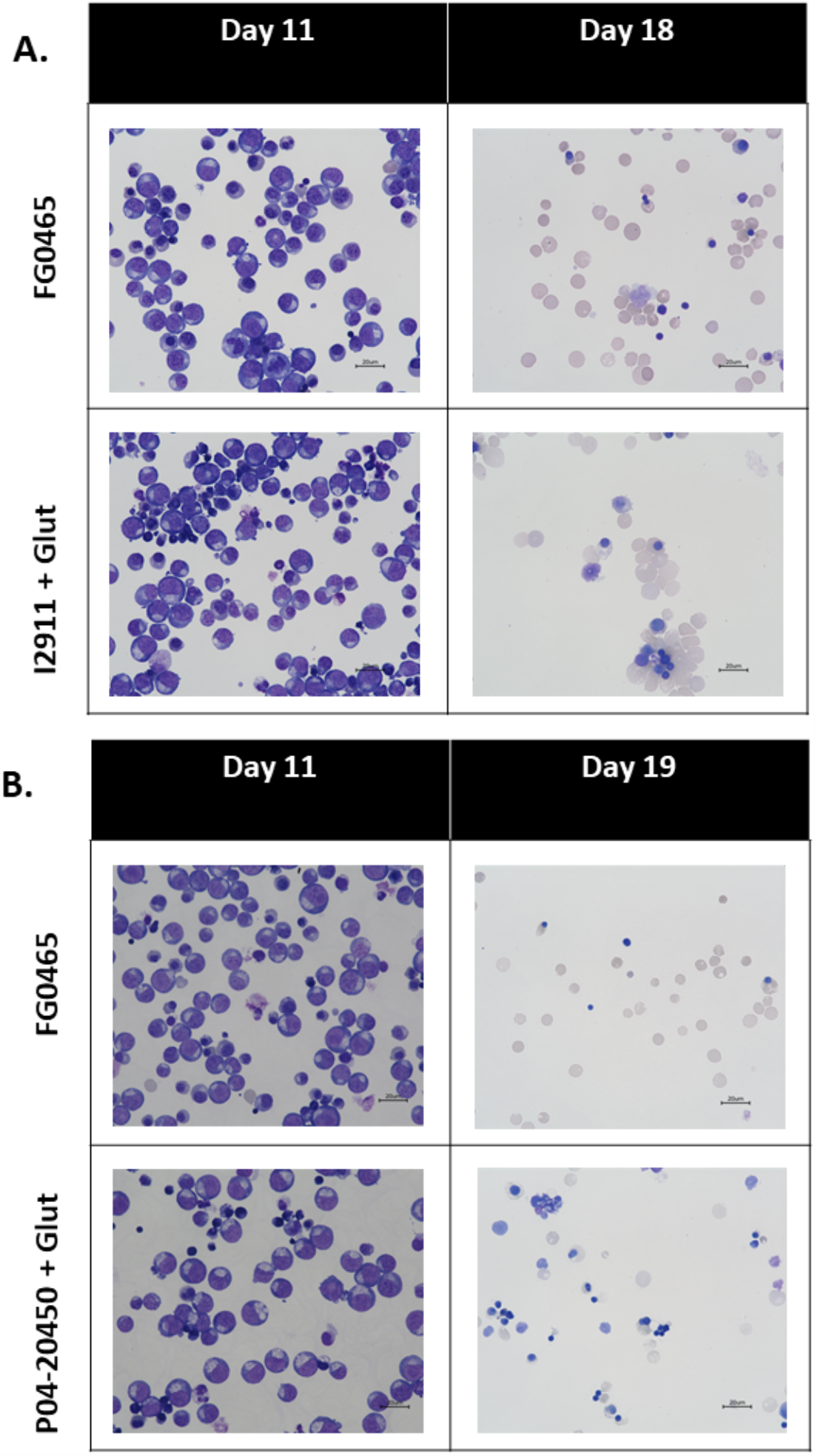
Differentiating erythroid cells grown in EDM made from different IMDM are morphologically similar. Representative images of May-Grünwald and Giemsa stained cytospins of: **(a)** day 11 and 18 cells grown in FG0465-EDM and I2911+Glut-EDM **(b)** day 12 and 19 cells grown in FG0465-EDM and P04-20450+Glut-EDM. Magnification, 60X; scale bar, 20μm.

To quantitate the enucleation rates, we used a nuclear dye and flow cytometry to distinguish between nucleated and enucleated cRBCs on day 18-19 (Fig. 3A). The results showed that 93.767% ± 0.895 (n=3) of cells grown in FG0465-EDM were enucleated, whereas only 75.35% ± 3.75 (n=2) of cells grown in I2911+Glut-EDM and 79.5% (n=1) of cells grown in P04-20450+Glut-EDM were enucleated (Fig. 3B). These results indicate that erythroid enucleation is influenced by the base IMDM media, with FG0465 promoting the highest enucleation efficiency.

**Figure 3:**
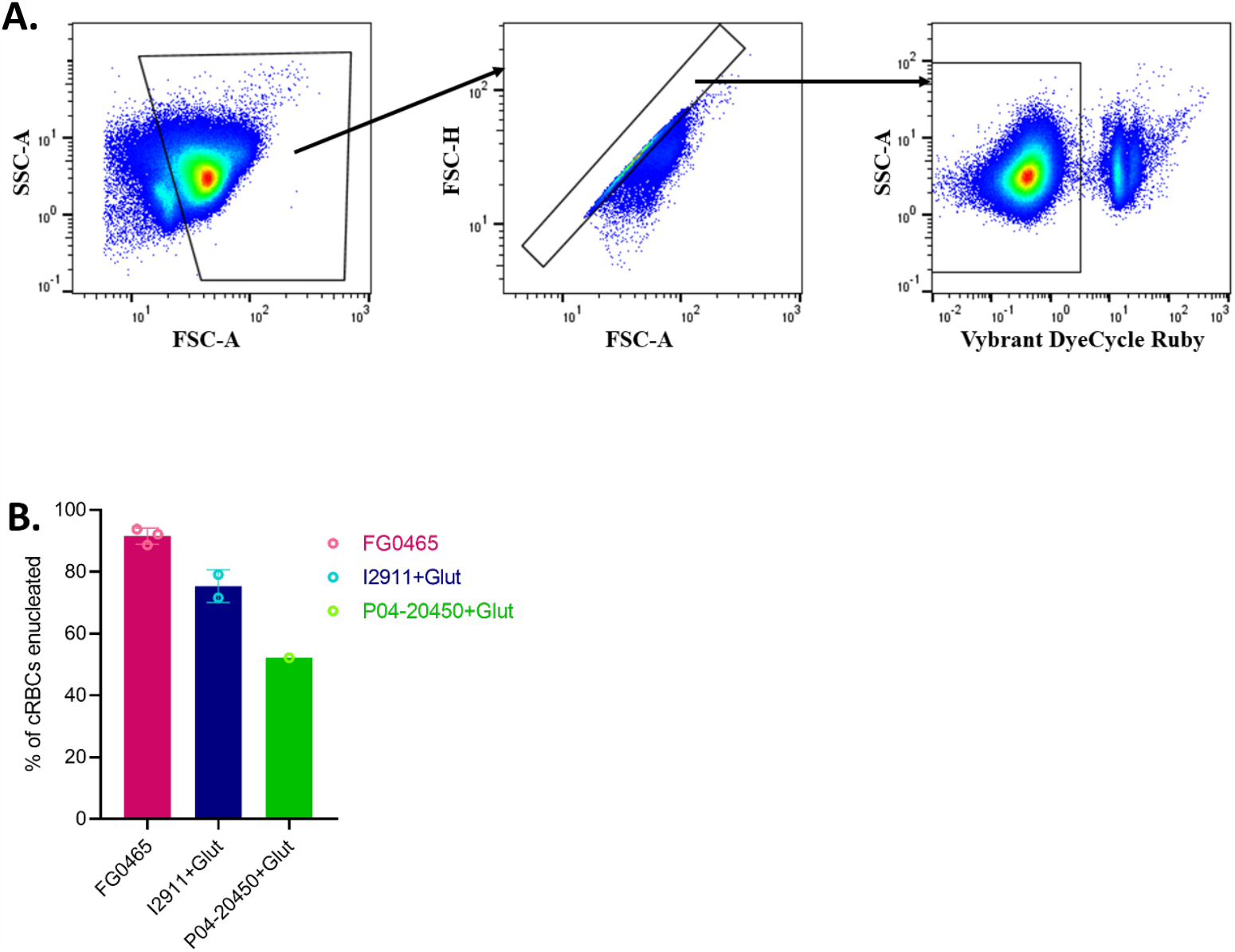
Enucleation occurs more efficiently in FG0465-EDM than in I2911+Glut and P04-20450 +Glut EDM. **(a)** Gating strategy (excluding free nuclei and restricting analysis to single cells). **(b)** Percentage of enucleated cRBCs assessed by flow cytometric analysis of Vybrant DyeCycle ruby nuclear dye-stained cRBCs on day 18 or day 19 of differentiation. Bars show mean +/-std. FG0465 (n= 3), I2911+ Glut (n=2), P04-20450 + Glut (n=1)

To further investigate how media formulations may differentially impact erythroid differentiation, we used a panel of antibodies to the surface proteins GYPA, CD49d, and CD71. which have well-defined patterns of expression during erythroid differentiation^13,14^. Using flow cytometry, we found that cell surface expression of these markers over the course of erythropoiesis was similar in the different EDM formulations, and followed the expected patterns for ex-vivo erythropoiesis when assessed individually. Specifically, the surface expression of GYPA remained high from day 12 to 17, the levels of CD71 started to decrease between day 12 and 17, and CD49d surface expression was detectable on day 12 but absent by day 17. One notable exception was an unexpected population of GYPA-null cells on day 17 in the P04-20450+glut-EDM (Fig. 4A and 5A). However, when CD71 and GYPA were analyzed together on double-stained erythroid precursor cells, we observed distinct patterns for cells grown in FG0465-EDM versus those grown in P04-20450+glut-EDM. (Fig.5A). Day 17 cells grown in P04-20450 EDM appeared intermediate between day 12 and day 17 cells grown in FG0465, with more cells expressing higher levels of CD71 than in FG0465 EDM (Fig. 5A). This suggests that the development of these cells may lag behind those grown in FG0465-EDM. This is further corroborated by the continued increase in enucleation for cells grown in P04-20450 EDM when cultured for longer as shown in Fig. 5B, from 52.2% on day 17 (compared to 88.7% for FG0465) to 85.2% on day 23 (compared to 98.6% for FG0465).

**Figure 4:**
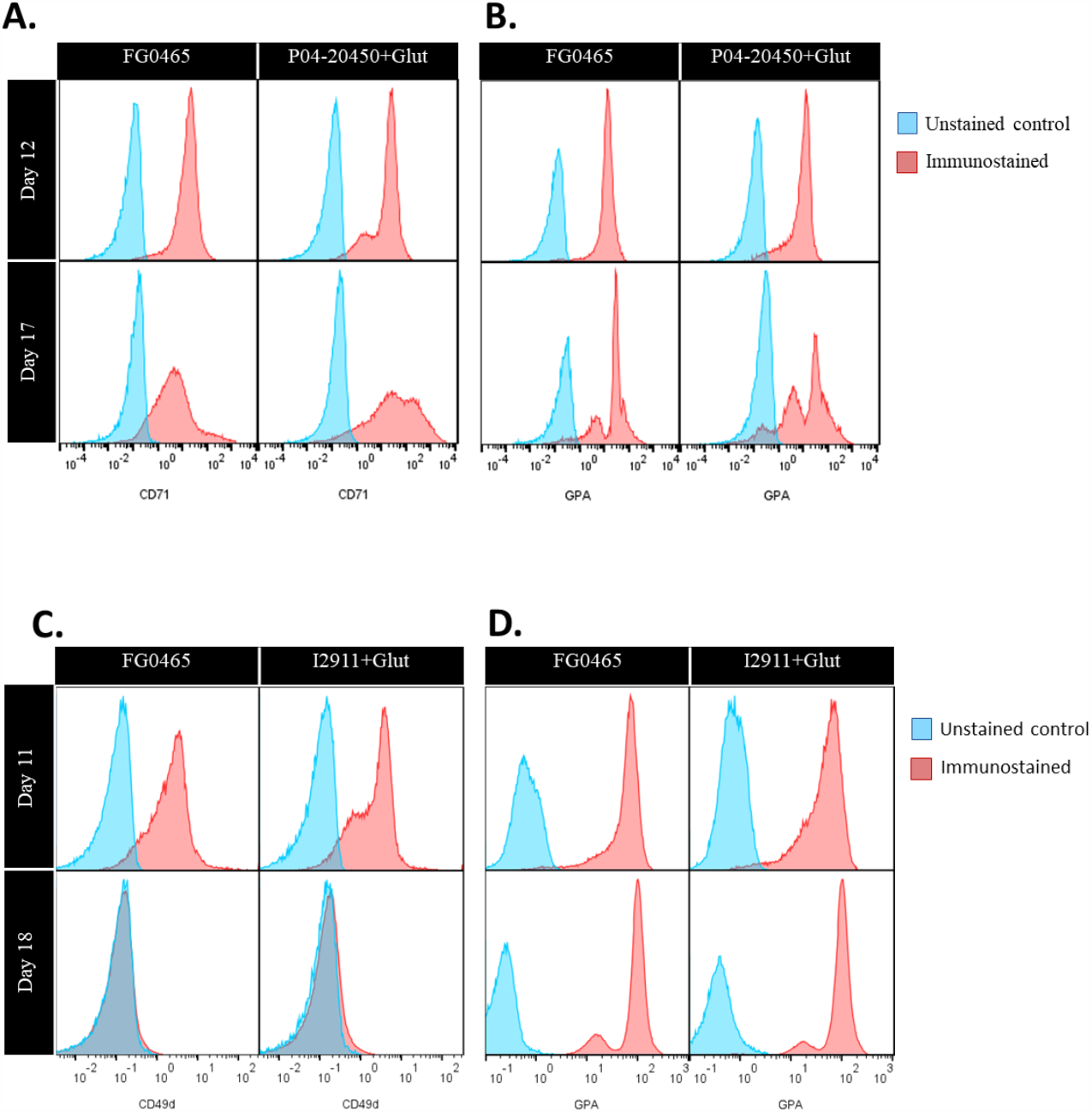
Expression of cell surface markers on erythroid cells grown in EDM made from the different IMDM formulations appear similar. Flow cytometric analyses of cell surface markers on cRBCs grown in FG0465, I2911+ Glut and P04-20450 EDM: **(a)** comparing CD71 surface expression on cells grown in FG0465 vs P04-20450 + Glut EDM on days 12 and 17 of differentiation **(b)** comparing GPA surface expression on cells grown in FG0465 vs P04-20450+Glut EDM on days 12 and 17 of differentiation **(c)** comparing CD49d surface expression on cells grown in FG0465 vs I2911+Glut EDM on days 11 and 18 of differentiation **(d)** comparing GPA surface expression on cells grown in FG0465 vs I2911+Glut EDM on days 11 and 18 of differentiation; all histograms normalized to unit area.

**Figure 5:**
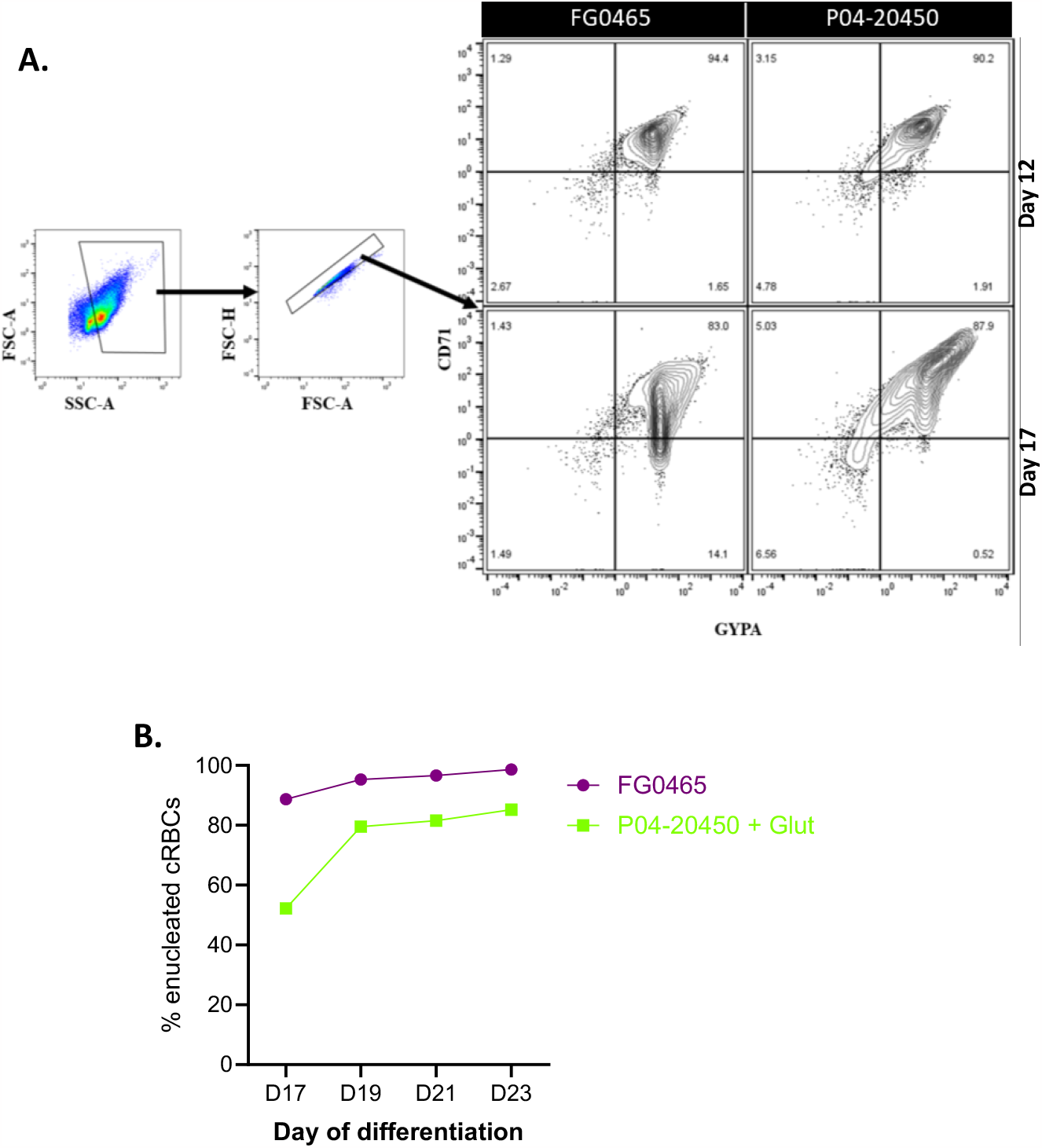
Differentiation of erythroid cells grown in P04-20450 EDM may lag behind FG0465 EDM-grown cells. **(a)** Comparing erythroid cell populations between FG0465 and P04-20450 EDM on days 12 and 17 by concurrently assessing expression of GPA and CD71 on doubly stained cells (showing gating strategy to exclude free nuclei and restrict analysis to single cells). **(b)** Comparing enucleation of cells grown in FG0465 and P04-20450 EDM from D17 to D23 of differentiation, assessed by flow cytometric analyses of vybrant-ruby nuclear dye-stained cells.

### Despite marketing similarities, published IMDM formulations differ in composition

Given the differences we observed in the growth rate and differentiation potential for primary human HSPCS grown in different EDM formulations, we requested the precise formulation sheets for each product for comparative analysis. EP-CM-L0216, FG0465, I2911 and P04-20450 are all described as IMDM containing sodium bicarbonate, glucose, phenol red and stable glutamine/ L-alanyl-L-glutamine, with P04-20450 described by Pan Biotech as equivalent to FG0465. EP-CM-L0216, I2911 and P04-20450 all reportedly contain cysteine, serine, tyrosine and valine, whereas FG0465 reportedly does not (Fig. 6A). All but EP-CM-L0216 are described as containing potassium nitrate (Fig. 6B). Other differences include slightly differing concentrations of disodium phosphate and sodium selenite (Fig. 6B), HEPES and phenol red (Fig. 5C), and riboflavin (Fig. 1D). The osmolality and pH reported by manufacturers for I2911 (276-304 mOS/kg, pH 7-7.4) and FG0465 (276-304 mOS/kg, pH 7-7.4) are similar. FG0465 is marketed as containing stable glutamine, and the product description page states that it contains L-alanyl-L-glutamine. However, it is notable that the formula information provided by Millipore Sigma upon request lists a concentration for L-glutamine, not L-alanyl-L-glutamine (Fig. 6A).

**Figure 6:**
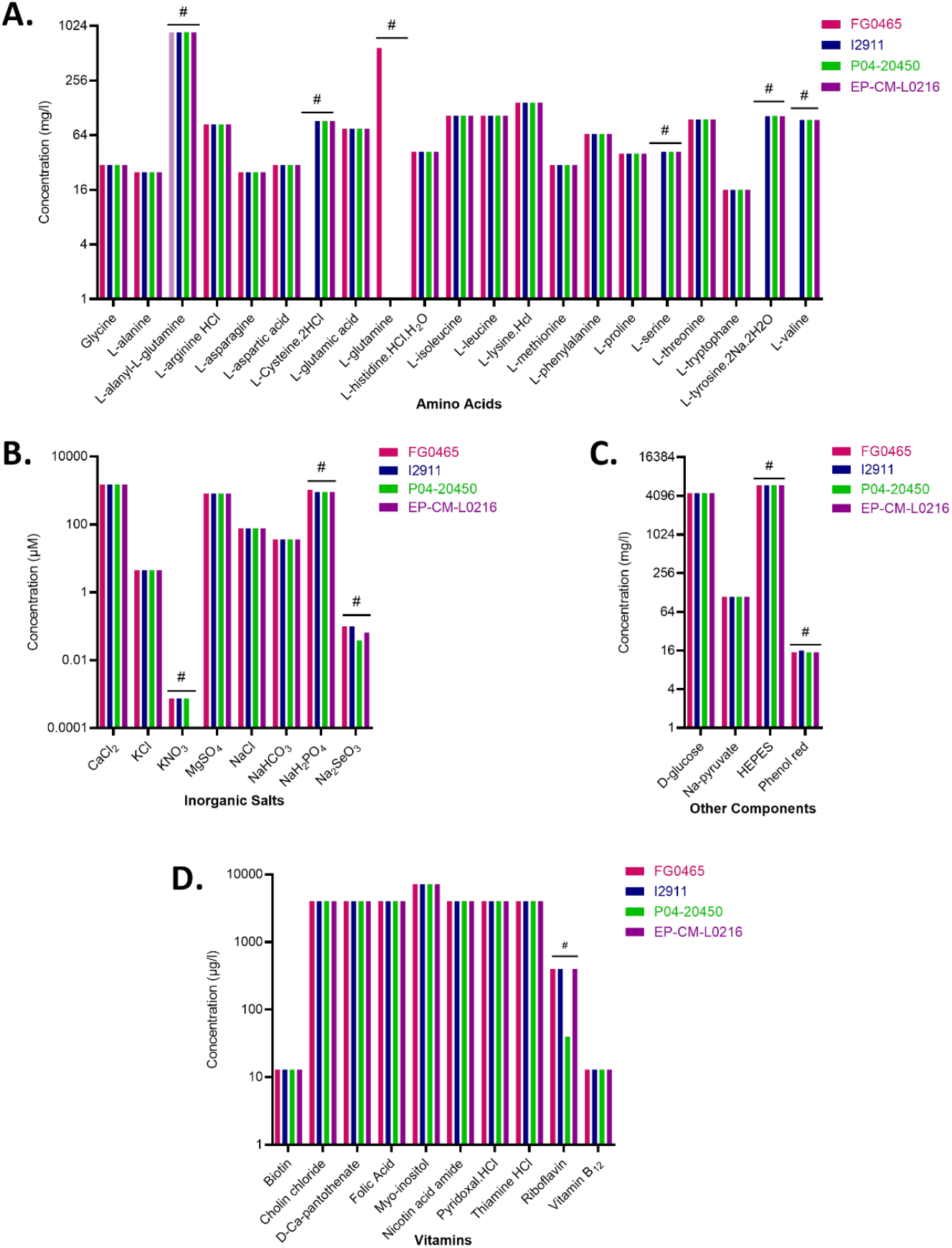
Manufacturers’ published formulation for IMDM products tested. **(a)** amino acid composition (**b)** inorganic salt composition (**c)** other components (**d)** vitamin composition. #, indicates where concentrations vary among the different products. All data shown in this graph is based on manufacturer’s published formulation except where grayed out if a modification is known to have been made but is not reflected on the manufacturer’s sheet.

## Discussion

The process of ex-vivo erythropoiesis of primary human HSPCs is resource-intensive; HSPCs and the associated reagents are expensive, and the procedure involves up to three weeks of cell culture at low density. Thus, it is important to maximize the yield of mature enucleated red blood cells that can be harvested at the end of this procedure for clinical or research applications. EDM, and therefore IMDM which is the basis of EDM, are integral to ex-vivo erythropoiesis. With different IMDM products marketed as being able to support stem cell cultures and the increasing frequency of supply chain issues impacting product availability, it is important to identify which formulation provides the optimum growth and differentiation media for erythroid precursors. In this paper, we have assessed the ability of four different IMDM products (FG0465, EP-CM-L0216, I2911 and P04-20450) to support the growth and differentiation of erythroid precursors into reticulocytes/ enucleated red blood cells. We have found that FG0465 yields higher erythroid cell expansion and supports greater levels of enucleation than all other products tested. I2911 and P04-20450 with additional glutamine were sufficient but not optimal for ex vivo erythropoiesis, while EP-CM-L0216 with or without added glutamine was not sufficient for ex-vivo erythropoiesis.

We found that IMDM products from different manufacturers with identical descriptions actually have different compositions. Do these differences in composition account for the differences in ability to support ex-vivo erythropoiesis? The one formulation we tested that could not support ex-vivo erythropoiesis at all was EP-CM-L0216. According to the published formulation data, EP-CM-L0216 is very similar in composition to I2911 and P04-20450, except that EP-CM-L0216 contains no potassium nitrate. To our knowledge, the importance of nitrates in ex vivo erythropoiesis has not been investigated, though the original formulation by Iscove contained nitrates^6,7^. While it is not possible to make a firm conclusion, from our data we speculate that the omission of potassium nitrate from EP-CM-L0216 accounts for the observed inability of this medium to support ex vivo erythropoiesis, as compared to FG0465, I2911+glut or P04-20450+glut. It is important to note that since we did not investigate the media composition beyond our reliance on the manufacturer’s published composition, we cannot rule out the possibility that other differences exist between the formulations that may account for the observed results.

Since erythroid precursors cultured in I2911+Glut-EDM and P04-20450+Glut-EDM are grossly similar to those grown in FG0465-EDM, our finding that the enucleation rates differed between these products and FG0465 was unexpected. One hypothesis is that these IMDM products do not support enucleation as well as FG0465. However, our data indicate that erythroid development in P04-20450+Glut EDM may lag behind erythroid development in FG0465 (Fig. 4A and 5A). With more time in culture, cells grown in P04-20450+Glut EDM can continue to enucleate to some degree. Since enucleated cells can be separated from nucleated precursors by some methods, including filtration or fluorescence-activated cell sorting following staining with a nuclear dye, P04-20450 and I2911 may still be useful for applications requiring enucleated red blood cells. However, the lower cell expansion observed in these IMDM products relative to FG0465 make these less attractive as alternatives to FG0465. In addition, the observation of a GYPA-null population on D17 in the cells grown in P04-20450+Glut EDM suggests this media may not be ideally supportive of erythroid development even beyond the effect on enucleation.

Ultimately, direct chemical analysis of the amino acid composition of EDM made from FG0465 and the other products tested here would reveal whether the final amino acid concentrations differ significantly among EDM made from these different products. Conversely, cystine, serine, tyrosine and valine could be added to FG0465, and potassium nitrate to EP-CM-L0216, to assess whether these changes lead to the phenotypes we would expect. These further experiments would be useful to understand the metabolite requirements of differentiating erythroid precursors. Altogether, this work has identified FG0465 to be superior to other IMDM products tested in ex vivo erythropoiesis. Many questions remain about why this is so, and this highlights the need to understand how individual media components affect ex vivo erythropoiesis.

## Acknowledgements

The authors thank Marilou Tetard and other members of the Egan Lab for helpful discussions. This work was supported in part by NIH DP2HL13718601, NIH 1R01HL166249-01A1, and a faculty scholar award from the Stanford Maternal Child Health Research Institute. E.S.E. is a Chan Zuckerberg Biohub San Francisco investigator.

## Author contributions

Conceptualization, data analysis, and writing, N.A.P. and E.S.E.; Performed experiments, N.A.P. ; Supervision and funding acquisition, E.S.E.

